# Deciphering Bacterial and Archaeal Transcriptional Dark Matter and Its Architectural Complexity

**DOI:** 10.1101/2024.04.02.587803

**Authors:** John S. A. Mattick, Robin E. Bromley, Kaylee J. Watson, Ricky S. Adkins, Christopher I. Holt, Jarrett F. Lebov, Benjamin C. Sparklin, Tyonna S. Tyson, David A. Rasko, Julie C. Dunning Hotopp

**Affiliations:** Institute for Genome Sciences, University of Maryland School of Medicine, Baltimore, MD 21201, USA; Department of Microbiology and Immunology, University of Maryland School of Medicine, Baltimore, MD 21201, USA; Center for Pathogen Research, University of Maryland School of Medicine, Baltimore, MD 21201, USA; Department of Microbial Pathogenesis, University of Maryland School of Dentistry, Baltimore, MD 21201, USA; Greenebaum Cancer Center, University of Maryland School of Medicine, Baltimore, MD 21201, USA

**Keywords:** transcriptomics, bacterial transcripts, archael transcripts, small RNAs, non-coding RNA (ncRNA), direct RNA sequencing

## Abstract

Transcripts are potential therapeutic targets, yet bacterial transcripts remain biological dark matter with uncharacterized biodiversity. We developed and applied an algorithm to predict transcripts for *Escherichia coli* K12 and E2348/69 strains (Bacteria:gamma-Proteobacteria) with newly generated ONT direct RNA sequencing data while predicting transcripts for *Listeria monocytogenes* strains Scott A and RO15 (Bacteria:Firmicute), *Pseudomonas aeruginosa* strains SG17M and NN2 strains (Bacteria:gamma-Proteobacteria), and *Haloferax volcanii* (Archaea:Halobacteria) using publicly available data. From >5 million *E. coli* K12 ONT direct RNA sequencing reads, 2,484 mRNAs are predicted and contain more than half of the predicted *E. coli* proteins. While the number of predicted transcripts varied by strain based on the amount of sequence data used for the predictions, across all strains examined, the average size of the predicted mRNAs is 1.6-1.7 kbp while the median size of the predicted bacterial 5’-and 3’-UTRs are 30-90 bp. Given the lack of bacterial and archaeal transcript annotation, most predictions are of novel transcripts, but we also predicted many previously characterized mRNAs and ncRNAs, including post-transcriptionally generated transcripts and small RNAs associated with pathogenesis in the *E. coli* E2348/69 *LEE* pathogenicity islands. We predicted small transcripts in the 100-200 bp range as well as >10 kbp transcripts for all strains, with the longest transcript for two of the seven strains being the *nuo* operon transcript, and for another two strains it was a phage/prophage transcript. This quick, easy, inexpensive, and reproducible method will facilitate the presentation of operons, transcripts, and UTR predictions alongside CDS and protein predictions in bacterial genome annotation as important resources for the research community.

## Importance

Our understanding of bacterial and archaeal genes and genomes is largely focused on proteins since there have only been limited efforts to describe the bacterial/archaeal RNA diversity. This contrasts with studies on the human genome, where transcripts were sequenced first through large scale EST sequencing projects that preceded the release of the human genome over two decades ago. We developed an algorithm for the quick, easy, inexpensive, and reproducible prediction of bacterial and archaeal transcripts from ONT direct RNA sequencing data. These predictions are urgently needed for more accurate studies examining bacterial/archaeal gene regulation, including regulation of virulence factors, and for the development of novel RNA-based therapeutics and diagnostics to combat bacterial pathogens, like those with extreme antimicrobial resistance.

## Introduction

Genomics, genome-enabled technologies, computational biology, and large-scale data mining are essential for rigorous, modern experiments on all organisms. Whole genome sequencing and protein-based annotation are now routine, low-cost approaches for analyzing bacteria and archaea. But often the annotation, and thus analysis and experimental validation, is limited to predicted protein-coding regions and a few highly conserved non-coding RNAs (ncRNAs) like the rRNAs. Yet, pathogen RNA transcripts, particularly ncRNAs and RNA-mediated regulation, offer an unexplored set of druggable targets, diagnostics, and potential therapeutics (1). In this context, a transcript is a physical RNA molecule that can be detected by sequencing RNA that has discrete start and end sites generated by a diversity of molecular mechanisms (*e.g.,* promoter/terminator, post-transcriptional processing). Transcripts are encoded within operons but are distinct from operons, which also include regulatory regions. Operons are widespread in bacterial/archaeal genomes, with ∼630-700 operons in *Escherichia coli* (2). Experimentalists have predicted operons using FPKM and/or sequencing depth without algorithms (e.g. (3, 4)), and efforts have been made to develop algorithms for their prediction (5–11). For example, the most recent version of Rockhopper predicts operons using a naïve Bayes classifier to combine strand, intergenic distance, and coordinated differential expression in a unified probabilistic model (12). Most operon predictions rely on the decades-old paradigm of operons as put forth by Jacob and Monod (13), which was summarized recently as “sets of contiguous and functionally related genes cotranscribed from a single promoter up to a single terminator” (14), including the operator regulatory region.

Fundamentally, the classical definition of operon is a DNA-based definition, defining a region in DNA that extends beyond the RNA-based transcripts to include the promoter/operator and terminator. Operons can have multiple transcripts due to post-transcriptional processing (15), alternate terminators (e.g. attenuation) (8, 16, 17), and alternate transcriptional initiation sites (14). There is a need for both DNA-based annotation of operons and RNA-based annotation of transcripts. Fundamentally, RNA-seq is transcript quantification, therefore it should be measured at the RNA/transcript level. Rockhopper has been used for differential expression of its predicted operons (9), but it yields different results than the corresponding transcript-focused analysis (14).

Currently mostly bacterial/archaeal RNA-seq studies are conducted using coding sequence (CDS) predictions. Even when issues with counting algorithms are mitigated for a CDS-focused analysis of polycistronic transcripts (18), measurements of CDSs in polycistronic transcripts are dependent on one another yet are treated as independent measurements with the statistics used to detect differential expression. This results in errors in variance estimations in differential expression tools (19). Comparisons of StringTie and Rockhopper have previously noted some of these issues, as well as the need for long RNA sequence reads to resolve these problems (8).

*E. coli* K12 is a well-studied genome that has some transcript predictions (17, 20), anti-sense RNA characterization (21), and transcriptional start site and terminator predictions (17, 22–25), all of which are aggregated and manually curated in RegulonDB (26) and EcoCyc (27). But even for this well studied organism, reference GFF files lack transcript annotations, and it can be difficult, if not impossible, to ascertain and use transcript structures for a differential expression analysis. The current work done to characterize transcripts and transcriptional regulation in *E. coli* (e.g., (26)), while laudable and necessary, is not possible for more than a few microorganisms, yet there is immense bacterial biodiversity. Therefore, we sought to develop a fast, simple, rigorous, and reproducible method for identifying bacterial transcripts that can be widely applied and takes advantage of recent advances in RNA sequencing, including PacBio IsoSeq and Oxford Nanopore Technologies (ONT) direct RNA Sequencing (14, 28–30). Transcript predictions will enable differential expression analyses using transcripts such that the analyses can be expanded to include non-coding RNAs (ncRNAs) and also use the latest transcript-based differential expression analysis tools like Salmon (31) and Kallisto (32). Transcript predictions are also needed to inform consequences of genetic knock-in and knock-out experiments (*e.g.,* (33)), identify regulatory sequences (*e.g.*, (8, 16, 34)) and detect post-transcriptional processing (*e.g.,* (15, 35)). Recent studies (8, 28, 36) reveal a much more complex picture of bacterial transcripts with post-transcriptional processing and potentially multiple promoters and terminators, including transcripts beginning or ending in the middle of adjacent coding sequences due to the coding density (17).

In this study, we describe a quick, easy, inexpensive, and reproducible method for whole transcriptome sequencing and annotation using ONT direct RNA sequencing. We directly test the methods on the *E. coli* K12 and E2348/69 strains and then also apply the algorithm to existing public data for *Pseudomonas aeruginosa* strains SG17M and NN2 (37), *Listeria monocytogenes* strains Scott A and RO15 (38), and *Haloferax volcanii* (39). Ultimately, we envision genomes where operons, transcripts, and UTRs are all annotated alongside CDSs and proteins in GFF files.

## Results

### ONT direct RNA sequencing of *E. coli* transcripts

We generated ONT direct RNA sequencing data (**Figure 1**) from RNA isolated from *E. coli* K12 and the pathogenic *E. coli* E2348/69 (40) grown at 37 °C with aeration in LB and DMEM media (**Table 1, Table A1**), which are virulence gene inducing growth conditions (15, 41–44). *E. coli* K12 is a well-studied genome including previous transcript predictions (17, 20), anti-sense RNA characterization (21), and transcriptional start site and terminator predictions (17, 22–25), all of which are aggregated and manually curated in RegulonDB (26) and EcoCyc (27). The inclusion of *E. coli* E2348/69 allows us to interrogate operon predictions in a related but clinically-relevant Enteropathogenic *E. coli* (EPEC) strain with plasmids (40) that has pathogenesis-associated operons, which have had fine scale analysis of transcription (15, 44). We focused on using ONT direct RNA sequencing, where RNA is sequenced directly in the pore (**Figure 1K**) to predict bacterial transcripts because it does not have template switching (36). Additionally, ONT direct RNA sequencing data lack genomic DNA contamination since sequenced RNA and DNA have markedly different signals with RNA advancing through the pore more slowly and with a higher electrical current range than DNA (**Figure 1E**). This difference between RNA and DNA is seen in every RNA read as the sequencing transitions from the DNA-based adaptor to the RNA, but is used to eliminate DNA reads with high fidelity. Therefore, we are confident that every read we analyze arose from a transcript, which is tremendously powerful when considering alternative transcripts, anti-sense transcripts, and non-coding RNA (ncRNA) predictions.

**Figure 1.**
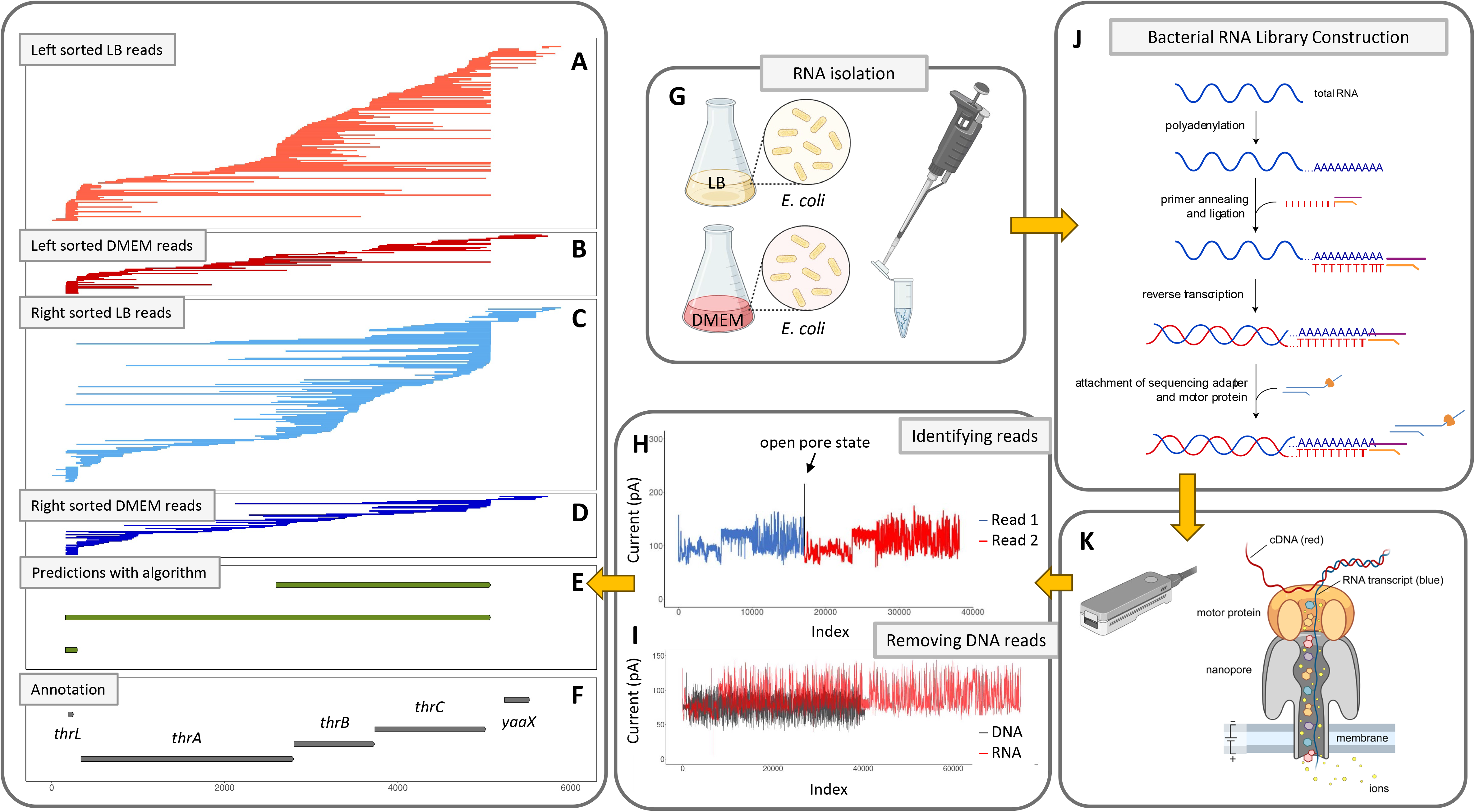
– Overview of the Experimental/Analysis Workflow. Plus-strand ONT direct RNA sequencing reads (shown as lines) are mapped from 1 bp to 6 kbp in the E. coli K12 genome (NC_000913.3), which corresponds to the thr operon, and sorted by their transcription stop site for E. coli K12 grown in rich LB media (left sorted, A; right sorted, C) and DMEM media (left sorted, B; right sorted, D). Our algorithm predicts 3 transcripts (E), and 4 CDSs in the GFF file are illustrated (F). The transcript for the leader peptide thrL is recovered in both growth conditions. (G) RNA was isolated from E. coli K12 grown at 37 °C with aeration in LB and DMEM media. (H) Squiggle plot for two sequencing reads in tandem. In this case, the open pore state was missed by the software resulting in a chimeric read. In both reads the DNA adapter can be observed with lower current followed by a relative flat plateau that corresponds to the polyA tail. This is followed by the current changes associated with the RNA moving through the pore. (I) Squiggle plots are shown of current for the same length DNA and RNA highlighting that the signal to base ratio is different for RNA and DNA. (J) The standard ONT direct RNA sequencing library was used on bacterial RNA that was in vitro polyadenylated following RNA isolation. Library construction and (K) loaded on an ONT MinION device for nanopore sequencing.

**Table 1.**
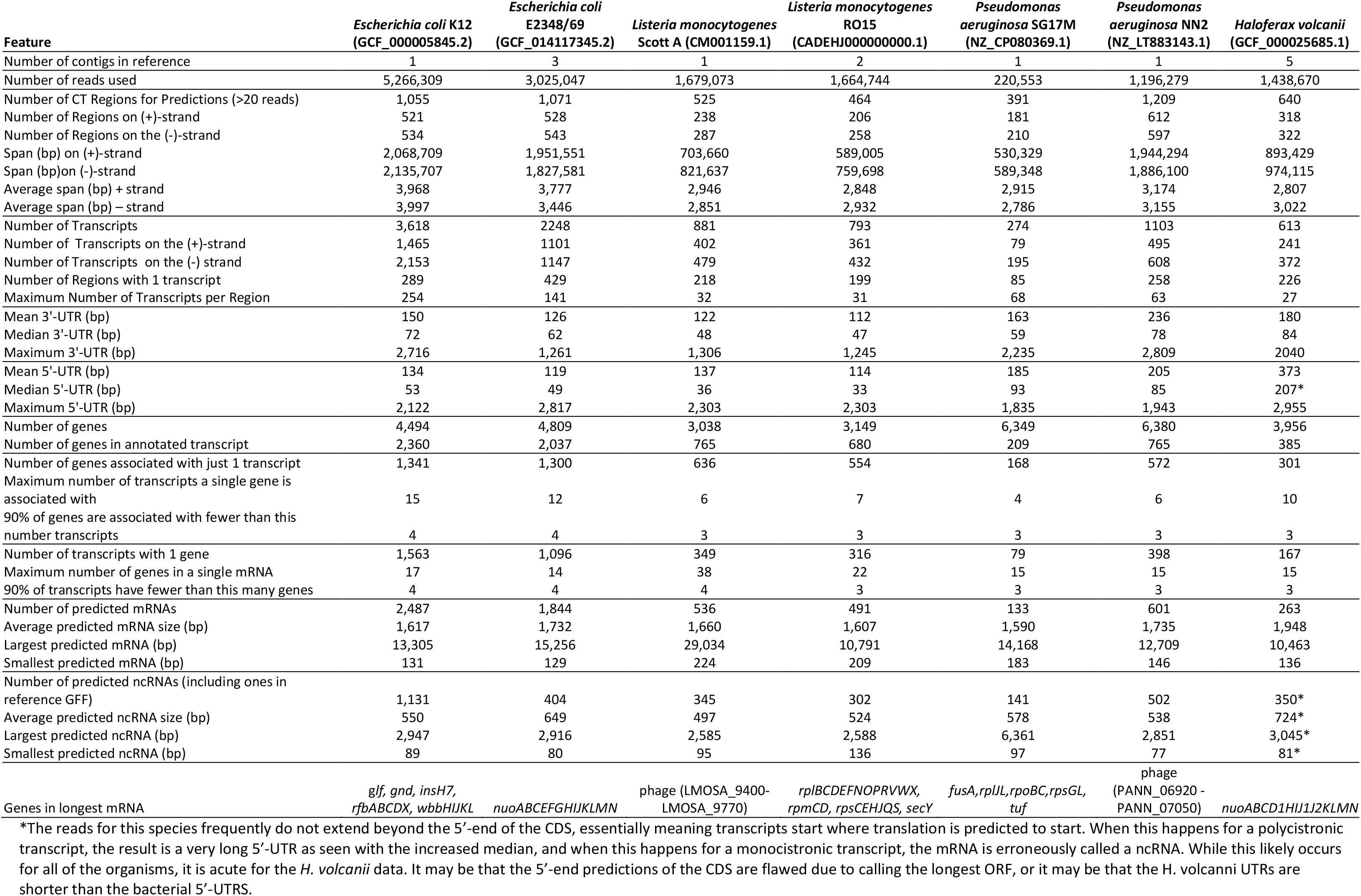
Characteristics of Predicted Transcripts for Escherichia coli, Listeria monocytogenes, and Pseudmonas aeruginosa.

### Predicted *E. coli* K12 transcripts

Using the 5,266,309 ONT reads generated for *E. coli* K12 (**Table 1**), we predicted transcripts using an algorithm we developed, which is described below. We identified 3,902 strand-specific contiguously transcribed (CT) regions in the K12 genome with 1,055 that have >20 reads that we used for predictions (**Table 1**). The 1,055 CT regions used for predictions are on average 4 kbp and include 521 regions on the (+)-strand spanning 2.07 Mbp and 534 regions on the (-)-strand spanning 2.14 Mbp (**Table 1**). There are 3,618 predicted transcripts with 1,465 predicted transcripts on the (+)-strand and 2,153 predicted transcripts on the (–)-strand (**Table 1**). There are 289 (27%) regions with only a single transcript predicted (**Table 1**), meaning the majority of CT regions contain more than one transcript either because operons overlap or because there are multiple overlapping transcripts.

Of the 3,618 predicted transcripts, 2,484 are predicted to be mRNAs and 1,134 are predicted to be ncRNAs (**Table 1**). mRNAs were defined as transcripts that have at least one annotated CDS found completely within the transcript boundaries, whereas a ncRNA was defined as a transcript that lacks a CDS found completely within the transcript boundaries. It is important to note that frequently the 5’-end of CDSs (and the N-terminal portion of the protein encoded by them) are incorrectly annotated, such that the assignment of transcripts as mRNA/ncRNA needs manual refinement in the future including possible curation of the N-termini of proteins; additionally, protein annotation may be informed and improved through transcript structural annotation. However, given these definitions, the average mRNA was 1,618 bp with the smallest and largest being 131 bp and 13,305 bp, respectively (**Table 1**). The average ncRNA was 517 bp with the smallest and largest being 52 bp and 2,947 bp, respectively (**Table 1**). Of these 1,134 predicted ncRNAs, 23 (2%) were already described in the reference GFF file and are ∼23% of the 98 previously annotated ncRNAs in the reference GFF file (**Table 1**).

Of the 4,494 annotated coding sequences (CDSs), 2,357 were in an annotated transcript while 2,775 were not, suggesting that with these growth conditions we could annotate transcripts for approximately half the CDSs. Of those, 1,341 (57%) CDSs were associated with a single transcript and 90% of CDSs were associated with <4 transcripts (**Table 1**, **Figure 2A**). While 1,564 of the predicted transcripts contained only a single CDS (**Table 1**, **Figure 2B**), the predicted transcript with the largest number of CDSs encoded within it contained 17 CDSs, including *glf*, *gnd*, *insH7*, *rfbABCDX*, and *wbbHIJKL* (**Table 1**). From the predicted mRNAs (excluding ncRNAs) and the predicted CDSs within those mRNAs, we predicted the 5’-and 3’-untranslated regions (UTRs). The median 5’-UTR is 53 bp and the most common length (mode) is 14 bp, while the median 3’-UTR has a median of 72 bp, and most common length (mode) of 36 bp (**Table 1**, **Figure 2CD**). There are concerns that ONT sequencing cannot capture the terminal 5’-end of transcripts. However, these results suggest that we are very close since it has been previously shown that the 5’-UTR is 20-40 nt (24).

**Figure 2.**
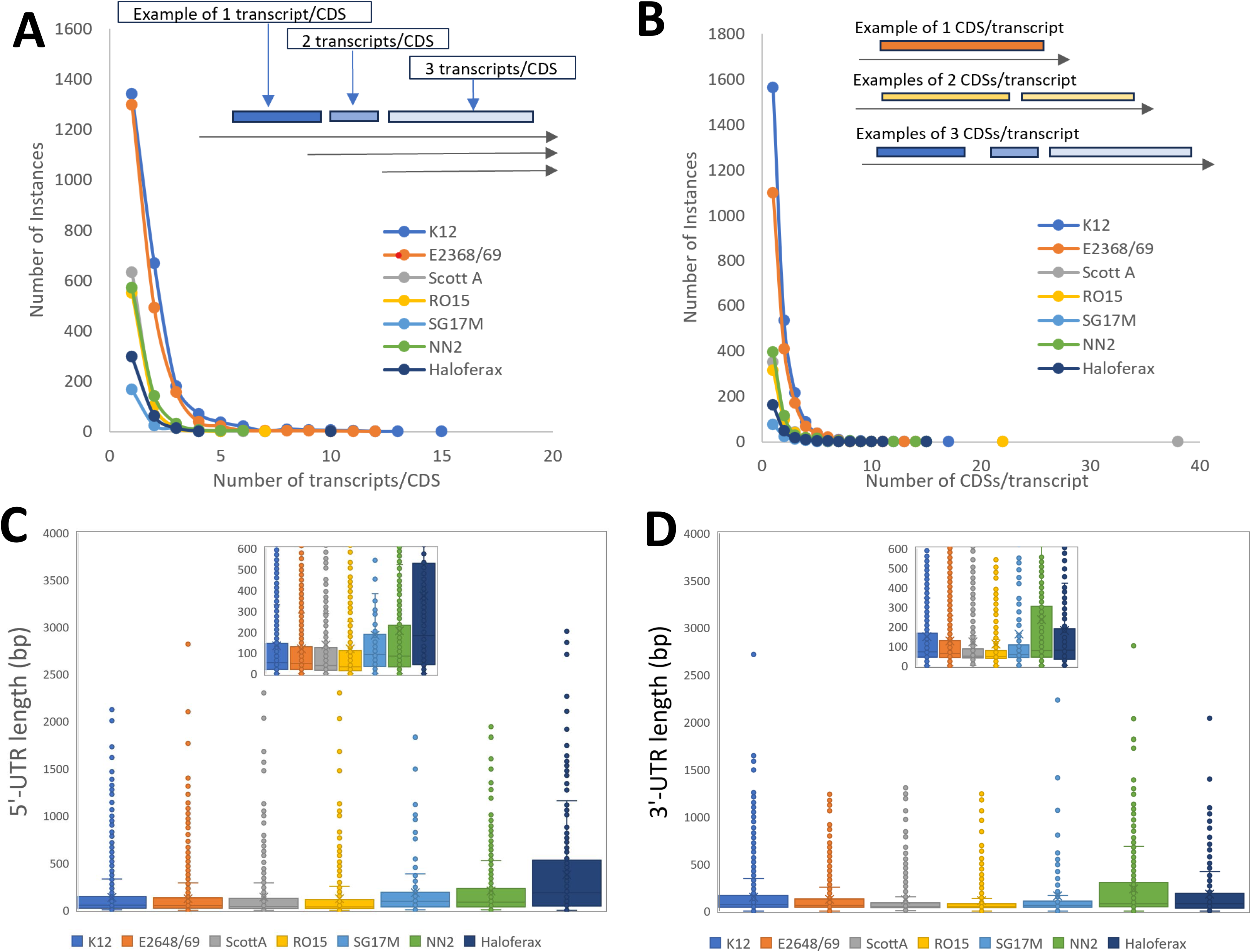
– Characteristics of Transcript Predictions. The distribution of the number of instances of CDS by transcripts/CDS (A) and the distribution of the number of instances of transcripts by CDSs/transcript (B) are shown for E. coli K12, E. coli E2368/69, L. monocytogenes ScottA, L. monocytogenes RO15, P. aeruginosa SG17M, P. aeruginosa NN2, and H. volcanii. The data points in these discrete distributions are connected by lines for visualization purposes. The inset in each illustrates how transcripts/CDS and CDSs/transcript are defined. The size distributions of predicted 5’-UTRs (C) and 3’-UTRs (D) are plotted for each of the six strains examined with an inset that zooms in on 0-350 bp to better illustrate the distribution of the majority of the data.

### Complexity of bacterial transcription

Our predictions detect tremendous bacterial transcript structural variation while confirming previous experimentally verified predictions. For example, in the *thr* operon, three transcripts are predicted, including the previously described *thrL* transcript for the leader peptide, the *thrLABC* transcript, and a *thrBC* transcript (45) (**Figure 1J**).

Other regions are more complex, like the region from 4,080-4,087 kbp encompassing *fdoGHI* and *fdhE* (**Figure 3**). EcoCyc and RegulonDB describe this entire region as an operon with two promoters—one that makes a transcript for the entire region and a second smaller internal transcript encoding *fdhE* that is started from a promoter within *fdoH* (**Figure 3**). The ONT data suggest differential expression of the transcript isoforms where *fdoGHI* are largely untranscribed in DMEM relative to LB while *fdhE* is transcribed in both (**Figure 3**). A small ncRNA is observed in DMEM when *fdoG* is not transcribed. (**Figure 3**). Our algorithm predicts 11 different transcripts in this entire region, including the *fdhE* transcript that starts in *fdoH* (**Figure 3**). As has been seen for decades in eukaryotic transcript prediction, automated predictions require manual curation. The algorithm likely underpredicts long transcripts, due to the limitations of the ONT technology as described below, so despite evidence for a complete *fdoGHI-fdhE* transcript, we do not predict it, likely because there is insufficient depth (**Figure 3**). But there is robust evidence for many of the other transcripts predicted that are not currently in RegulonDB, EcoCyc or the GFF file, including a transcript of just *fdoG*, just *fdoGHI*, two putative overlapping small RNAs that overlap the end of *fdoI* and the beginning of the *fdhE* transcript, and four putative overlapping small RNAs that overlap the beginning of *fdoG* (**Figure 3**). In a typical differential expression analysis that uses CDS regions., these four putative small RNAs overlapping *fdoG* would likely be misinterpreted as expression of *fdoG* in DMEM. Importantly, while we detect these transcripts, we cannot ascertain that they have a function, and they could merely be stable degradation products of transcription. Regardless, they are likely to confound and obfuscate differential expression analyses.

**Figure 3.**
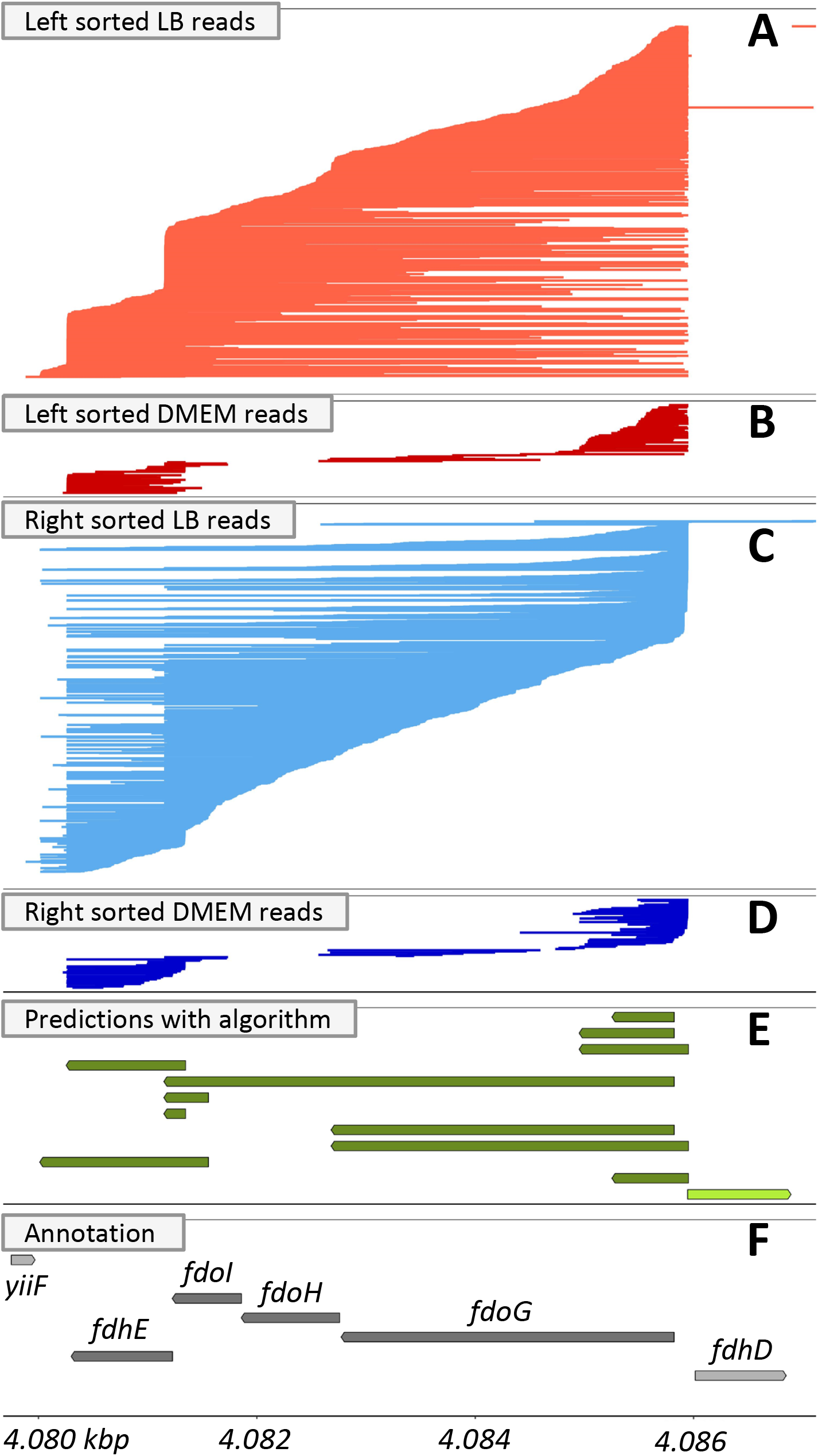
– fdoGHI-fdhE Transcripts. Reads mapping to the minus strand of the E. coli K12 genome (NC_000913.3) grown in LB (A, C) and DMEM (B, D) are shown for a region from 4,080-4,088 kbp. To facilitate the visualization of the starts and stops of transcripts, reads were sorted by either their left most (A, B) or right most (C, D) position and plotted from top to bottom accordingly. Transcript predictions from our algorithm (E) and the predicted CDSs in the reference gff file (F) are shown with arrows indicating the direction of transcription and with transcripts/CDSs on the different strands having different shading (light for the (+)-strand and dark for the (-)-strand).

Across the 11 transcripts predicted in the *fdoGHI/fdhE* region, there is imprecision in transcript start and end sites, as previously described (15, 24). This variability includes slightly longer transcripts that extend beyond *fdhE* that are observed under both growth conditions and was reproducible across all sequencing runs (**Figure 3**). This variability is seen in many regions, suggesting that transcription and termination are flexible.

### Predicted *E. coli* E2348/69 transcripts

The 60% fewer reads sequenced for *E. coli* E2348/69 relative to K12 led to fewer transcript predictions (**Table 1**), particularly fewer ncRNA predictions, but otherwise the results are quite similar. The longest predicted mRNA for E2348/69 is *nuoABCEFGHIJKLMN,* a known operon (46, 47). Unlike the K12 strain, the E2348/69 strain contains two plasmids (NZ_CP059841.1 and NZ_CP059842.2, respectively) and mRNA and ncRNAs were predicted on both plasmids. Of the 405 predicted ncRNAs, 3 (1%) were already described in 4 ncRNAs in the reference GFF. Additional known ncRNAs missing in the reference GFF file were identified, including *glmY* and *glmZ*, both of which are important for regulation of the *LEE* operon and thus virulence (44).

The transcription of *LEE* operons, which are found in the E2348/69 genome, has been extensively studied. In *LEE4,* a promoter upstream of *sepL* produces a *sepL-espADB* transcript that is post-transcriptionally cleaved with RNAse E to generate an *espADB* transcript and a *sepL* transcript that is then further endonucleolytically degraded (15) (**Figure 4**). A putative transcriptional terminator was previously identified downstream of *espB* within *cesD2*, but it was hypothesized that there is readthrough transcription of the terminator (15). The ONT sequencing data here provide evidence for readthrough of the transcriptional terminator. Very few reads included both the *cesD2-vapB-escF* region and *sepL*, which may be an indication that processing to remove *sepL* is more efficient on the longer transcript that terminates after *espF*, although we can’t rule out that the 6 kbp transcript of the whole region was not predicted due to the size limitations of ONT direct RNA sequencing. Consistent with the latter, the 4 kbp *sepL-espADB* transcript has been detected by Northern blots in multiple studies (15, 44), yet it is very infrequently detected here. Prior 5’-and 3’-RACE of LEE4 transcripts revealed variation in transcript ends, which we also detect, with multiple reads supporting a longer transcript at the 5’-end of *sepL,* which seems to be a frequent phenomenon across all transcripts. We also predict single CDS transcripts that encode for *espA*, *espB,* and *espF*.

**Figure 4.**
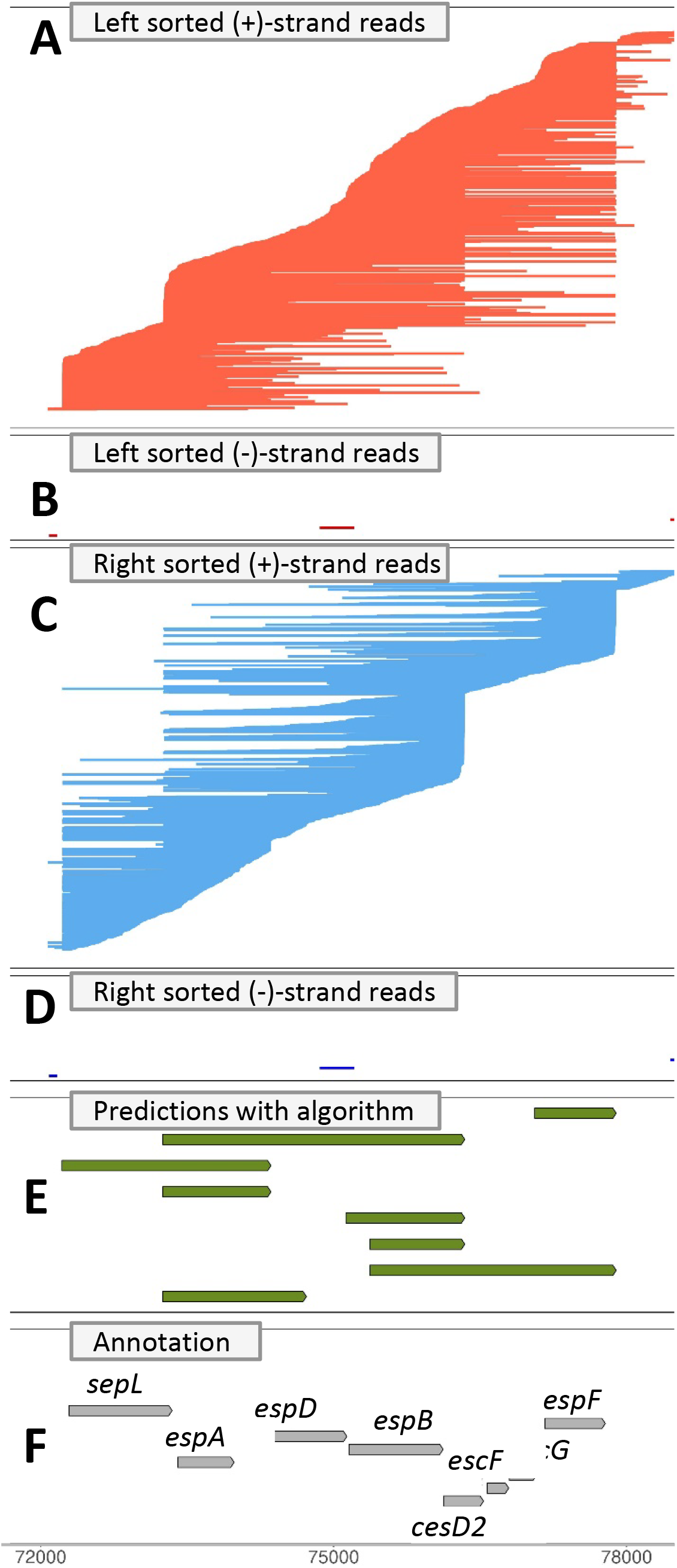
– LEE4 Operon. Reads are illustrated that map to the plus strand (A, C) and minus strand (B, D) of the E. coli E2348/69 genome (GCF_014117345.2) grown in LB or DMEM for a region from 72-78 kbp. There are no reads from the LB conditions on the (+)-strands. To facilitate the visualization of the starts and stops of transcripts, reads were sorted by either their left most (A, B) or right most (C, D) position and plotted from top to bottom accordingly. Transcript predictions from our algorithm (E) and the predicted CDSs in the reference gff file (F) are shown with arrows indicating the direction of transcription and with transcripts/CDSs on the different strands having different shading (light for the (+)-strand and dark for the (-)-strand).

### Data re-use and transcripts in Listeria monocytogenes, Pseudomonas aeruginosa, and Haloferax volcanii

Through data re-use, we also predicted transcripts using published ONT data for *P. aeruginosa* strains SG17M and NN2 (37), *L. monocytogenes* strains Scott A and RO15 (38), and *H. volcanii* (39). All five of these strains had fewer sequencing reads than we had for *E. coli*, leading to fewer predictions of transcripts, including both mRNA and ncRNA (**Table 1**). Yet we were still able to predict 274-1103 transcripts across the five strains and those transcripts were similar to the *E. coli* data with respect to mean/median/mode 3’-UTR lengths, proportion of single CDS transcripts, proportion of single transcript CDSs, size distribution of mRNA, and size distribution of ncRNA (**Table 1**). The 5’-UTR predictions were similar across the bacterial strains, but the archaeal reads frequently did not extend beyond the 5’-end of the CDS (**Table 1**). For a monocistronic transcript, the mRNA is erroneously called a ncRNA, while for a polycistronic transcript, a very long 5’-UTR is predicted resulting in an increased median (**Table 1**). It may be the 5’-end predictions of the CDS are flawed due to calling the longest ORF, or it may be that the *H. volcanni* UTRs are shorter than the bacterial 5’-UTRS and/or not well captured with the ONT technology. Across all seven strains examined, the longest transcript varied, although two were phage transcripts and two were *nuo* transcripts (**Table 1**). The inclusion of *L. monocytogenes* was an important test case since it is a firmicute with leading strand transcription bias (48), which led to fewer and longer CT regions, but did not prevent high quality transcript predictions. While there was ONT direct RNA data for further species of gamma-Proteobacteria, we limited this analysis to just two species with two strains each from this taxon. Overall, these results suggest that this simple sequencing method combined with our algorithm can be applied widely to archaeal/bacterial genomes to enable rigorous and robust transcript predictions.

### Differential expression analysis

These transcript predictions can be used in a differential expression analysis using Salmon and EdgeR as demonstrated with existing E2348/69 short read data from the SRA (PRJEB36845/E-MTAB-88804) and the long read ONT data generated here (**Figure 5**). Even when analyzing the same Illumina data, there is discordance between the TPM values calculated for transcripts and CDSs (**Figure 5GHI**). There is also discordance when quantifying the ONT data, which might be attributed to many factors, which bears further investigation but is beyond the scope of this manuscript. We have concerns about using ONT reads for differential expression analysis since shorter transcripts are preferentially sequenced relative to longer transcripts (**Figure 6F**, as described below). In addition, the larger numbers of Illumina reads generated is beneficial in the calculation of TPMs and subsequent statistical analyses.

**Figure 5.**
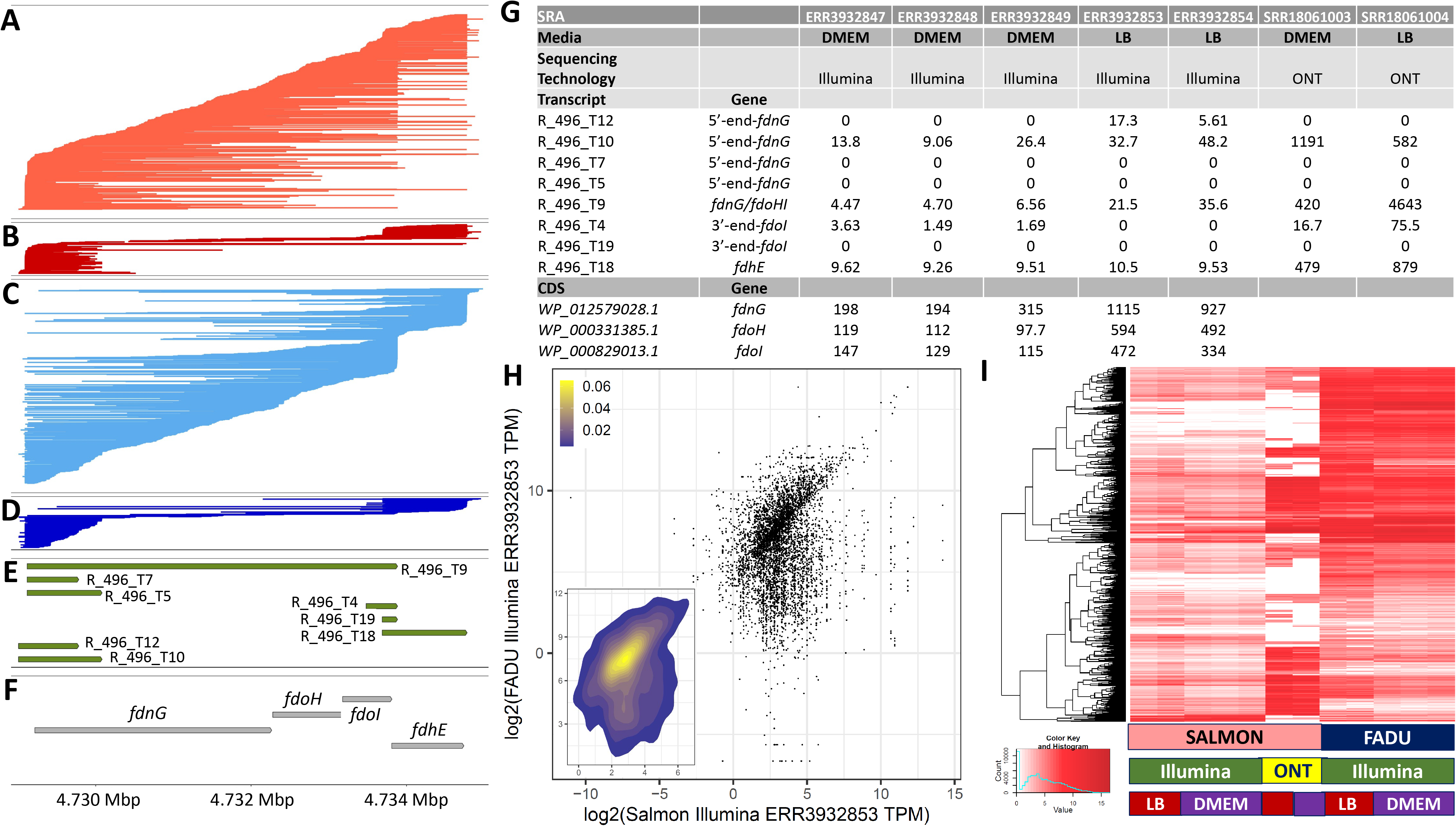
– Differential expression of predicted transcripts. Reads are illustrated mapping to the plus strand of the E. coli E2348/69 genome (GCF_014117345.2) grown in LB (A, C) or DMEM (B, D) from 4.730-4.735 Mbp sorted by either their left most (A, B) or right most (C, D) position. Transcript predictions from our algorithm (E) and the predicted CDSs in the reference gff file (F) are shown with arrows indicating the direction of transcription. Table of TPMs calculated with Salmon for transcripts and FADU for CDSs (G) for the same region shown in panels ABCDEF. For ONT reads, only Salmon was used. Plot of the log2(TPM) for all CDSs and all corresponding transcripts for ERR393285 (H). Heatmap clustered by genes for the log2(TPM) for all CDSs calculated with FADU and all corresponding transcripts calculated with Salmon for Illumina and ONT reads generated from LB and DMEM (I).

**Figure 6.**
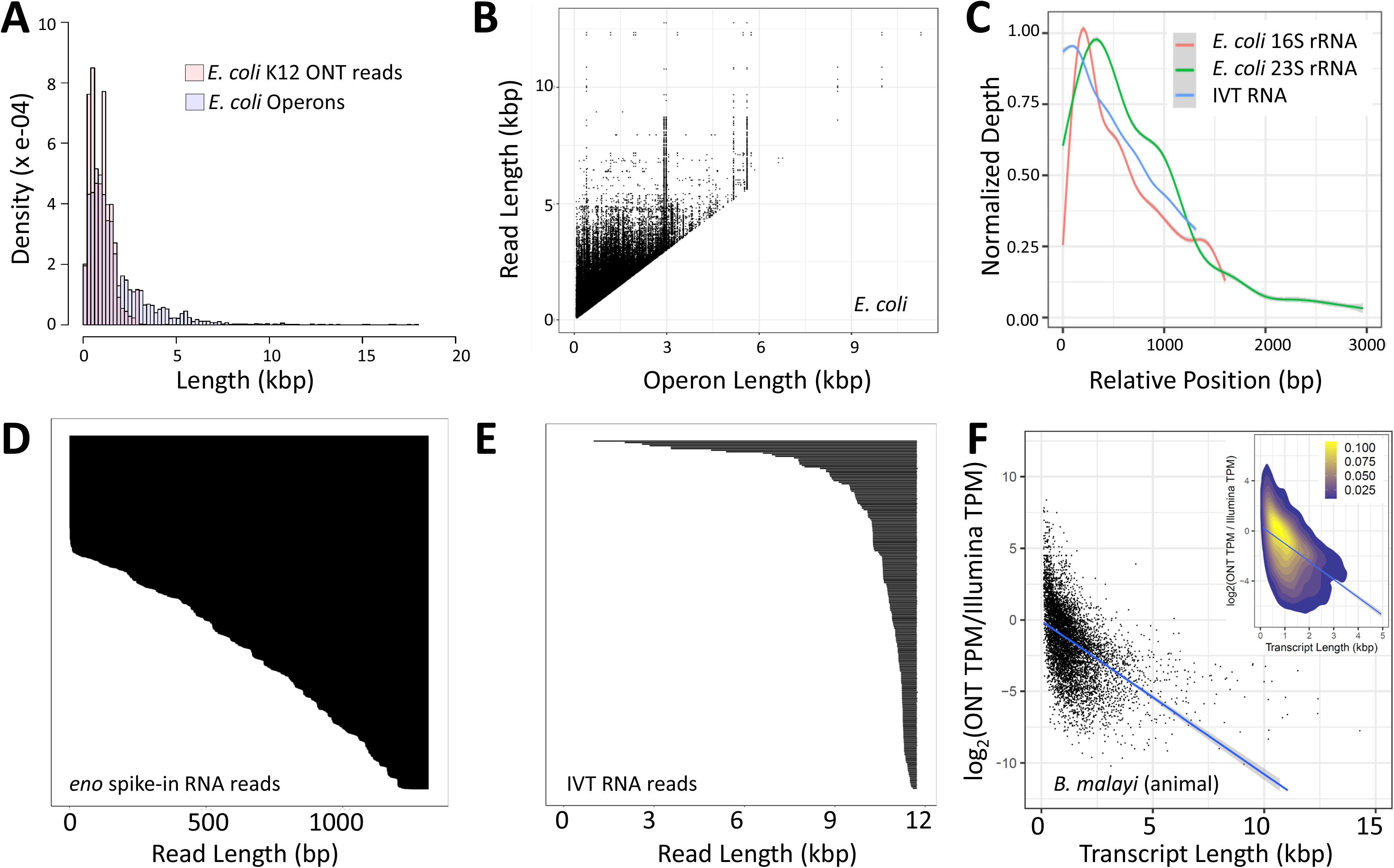
– ONT sequencing characteristics that informed algorithm development. Size distribution of all of the E. coli K12 ONT sequencing reads aligning outside the rRNA reads compared to the distribution of predicted operons (A). For the 285,619 reads that are longer than the operon they map to, the length of reads is plotted relative to the size of the operon they map to (B). Normalized sequencing depth from the 3’-end to the 5’-end for E. coli K12 16S rRNA, E. coli K12 23S rRNA, and IVT RNA (SRR23886069), all thought to be complete, showing the 3’-bias in sequencing (C). Distribution of read lengths for the 1.3 kbp yeast enolase ONT spike-in (D) and an 11.7 kbp IVT RNA (E) from SRR23886069where only reads ending at the far right position are shown. The log transformed ratios of Illumina (SRR3111494) and ONT (SRR23886071) TPM values for RNA isolated from adult female Brugia malayi, a filarial nematode and invertebrate animal, is compared to the transcript length, illustrating how shorter transcripts have more Illumina reads relative to ONT reads than longer transcripts (F). Our interpretation is that ONT sequencing is biased toward shorter transcripts. The inset uses the heat function to show the intensity of the points in the region which contains most of the data.

### Features and Limitations of ONT direct RNA sequencing of *E. coli* transcripts

To develop rigorous methods and algorithms to predict these transcripts, we needed to understand the characteristics of ONT direct RNA sequencing of bacterial transcripts, which we expect to differ from sequencing of eukaryotic transcripts given the differing physical features and stability of prokaryotic and eukaryotic RNA. Overall, operons >5 kbp are difficult to obtain in a single read (**Figure 6A**), but reads can be sequenced that span most predicted operons as well as exceed the boundaries of an existing operon prediction (**Figure 6AB**). While *E. coli* has known transcripts >10 kbp, we do not generate reads >9 kbp (**Table 1**). This is, at least in part, likely due to the ONT technology since we observe that (a) this is reproducible across multiple systems and RNA molecules we know must be full length, like rRNAs (**Figure 6C**), (b) there is 5’-truncation of transcripts in 11.7 kbp full-length *in vitro* transcribed (IVT) polyadenylated RNA (**Figure 6D**), and (c) there are many incomplete reads for the 1.4 kbp yeast enolase 2 (ENO2) RNA calibration strand provided by ONT (**Figure 6E**). Sequenced transcripts are also 3’-truncated (**Figures 1ABCD, 3AC, 4ABCD)**, as previously described for ONT (28, 36, 37) and PacBio IsoSeq (30) sequencing of bacterial transcripts, possibly from (a) random fragmentation of RNA, (b) RNA degradation, and/or (c) incomplete transcription in a bacterial cell. Additionally, we found that shorter transcripts are preferentially sequenced relative to longer transcripts (**Figure 6F**). This is despite counts/RPKMs being reported as well correlated between Illumina cDNA-based sequencing, ONT cDNA-based sequencing, and ONT direct RNA sequencing (49), as well as when nanopore direct RNA sequencing CPMs are compared to the absolute concentration of a spike-in (50).

To address incomplete reads and preferential sequencing of shorter transcripts, we developed a method that first predicts transcript start/stop sites in locations where there is an over-abundance of reads starting and ending. Subsequently, the actual transcripts are defined by measuring the strength of the connection between those start and stop sites using a model that supports the characteristics of truncated transcripts where smaller transcripts are preferentially sequenced. In this way, we can predict 12-15 kbp mRNAs (**Table 1**), despite having a shorter max ONT read length (**Figure A1**).

One of the features of ONT direct RNA sequencing is the ability to use changes in electrical current to detect RNA modifications including *N*6-methyladenosine (m^6^A), 5-methylcytosine (m^5^C), inosine, pseudouridine, and many more (51). At a minimum, posttranscriptional modifications are expected in bacterial tRNA and rRNA (52), but might also be present in mRNA and would lead to nonrandom changes in sequencing depth and base calling errors (53, 54). To alleviate this issue, we use a depth calculation computed assuming every base is equally present in a read using start/end positions of bed files for mapped reads. This also enables predictions in the presence of errors in the reference or sequence divergence from the reference (e.g. (55)).

Using only read end positions may also facilitate predictions of transcripts for one strain using data from a different strain. However, given that we haven’t ascertained how much transcript structural diversity there is between strains, it may be ill-advised. For that reason, we did not, for example, use the SG17M and NN2 data to make available predictions for the research community for the frequently used *P. aeruginosa* PA01.

Chimeric RNA sequencing reads were detected in all samples, including chimeras between the ONT ENO2 calibration strand and sample RNA (**Figure 1H**). A subset of these are *in silico* chimeric reads, with a spike observed in the electrical current when analyzing the raw signal data, indicating an open pore state that was missed by the MinKNOW software. Others lack this spike and could be either ligase-mediated chimeras or *in silico-*mediated chimeras where the open pore state was too short to be detected (**Figure A2**) (56). In our analysis, this was addressed by removing the clipped portions of mappedreads. When mapping reads to a reference genome, portions of a mapped read that do not align with the reference will be either “soft-clipped” or “hard-clipped.” A soft clipped read has a portion that does not align to any other area of the reference (e.g. the ENO2 portion of an ENO2/mRNA chimeric read), whereas a hard clipped read has two portions that align to different parts of the genome. For soft-and hard-clipped reads we used the primary alignment, ignoring the clipped portion of the read.

### Transcript Prediction Algorithm

We developed an algorithm to predict transcripts based on these characteristics and applied it as described above. Each CT region is examined separately along with the reads completely contained within that region. CT regions are initially defined through the bed input file and subsequently refined to subdivide regions based on a minimum depth cut-off (default=2). Ultimately a region needs to have a minimum number of reads fully contained within it to be considered (default=2). The change in depth of the sequencing reads for each genomic position of the CT region (*D*reg) ignoring mismatches/indels is calculated as

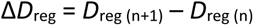

Potential start and stop sites are predicted at positions where Δ*D*reg is sufficiently positive/negative and always includes the first and last position of the region. These predictions require that |Δ*D*reg| surpasses a threshold (default=4). ONT sequencing has issues identifying precise ends of transcripts due to polyA-trimming as well as sequencing 5’-ends, such that predicted start/stop sites in close proximity (default=100) are grouped.

Candidate transcripts are predicted using the Cartesian product of all predicted start and stop sites. The total read count (Ntot) is calculated from the number of total reads that are mapped to all transcripts that fully contains them, allowing for mapping to multiple transcripts. The count of exclusively assigned reads (Nea) is calculated after mapping each read to the shortest transcript that fully contains it. The candidate transcripts are processed from shortest to longest computed as *Ratio* = Nea / Ntot. If this ratio is less than the threshold (default=0.2), the candidate transcript is discarded. If possible, reads from discarded transcripts are re-assigned to longer transcripts, and the Nea is recalculated such that reads initially assigned to now discarded transcripts can be used to support a longer transcript. All transcripts that meet the ratio at the end of the analysis are reported in a gff file and a bed file.

### Assembling ONT RNA reads

We attempted to assemble the ONT direct RNA reads with existing tools, including TAMA (tc_version_date_2020_12_14) (57), Cupcake (v.29.0.0) (58), and StringTie (v1.3.4d) (59). None of the existing tools recapitulated the complexity of the bacterial transcripts accurately, leading us to develop a new algorithm for the prediction of bacterial transcripts (**Figure A3**).

## Discussion

In biology, dark matter is often used to describe functional portions of genomes that are not described or annotated. In most bacteria, transcripts are largely dark matter, with CDSs often serving as a proxy, albeit a poor one. Here, we show that bacterial long read transcriptome data can be used to predict bacterial transcripts using a tool designed for the complexities and nuances of prokaryotic transcripts. Application of this tool to ONT data from three organisms revealed extensive transcript structural variation, transcription of RNA on both strands in some regions, overlapping transcripts, and a diversity of non-coding RNAs. Fundamental biological differences such as a high coding density and polycistronic transcripts in bacterial genetics lead to problems in applying transcript prediction tools developed for the human genome. We cannot merely apply the same laboratory and computational methods that were designed and optimized for humans and eukaryotic model organisms, with the false assumption that they will work because bacteria are “simpler” than humans.

The transcript structural diversity highlights the need for algorithmic and analysis improvements that are important for rigorous differential expression analyses, molecular evolution analyses, and other analyses as well as laboratory experiments like making knock-outs/ins or promoter analysis. Coupling this with a re-analysis of existing *E. coli* proteomics data would be enlightening and informative in understanding if transcripts annotated as ncRNAs are producing previously undescribed proteins/peptides.

Yet, there is much room for improvement for bacterial transcript predictions, both through lab experimentation and bioinformatics. We hope that attention to the recent developments in bacterial transcript sequencing will lead to the development of more bioinformatics tools with a richness like that seen for eukaryotic transcript prediction. The greatest improvement in the lab would be in obtaining more full-length reads, particularly for long transcripts, which is a challenge for all long-read sequencing platforms. For ONT, the new chemistry may improve the length, and further improvements may be possible by altering the reverse transcription method needed to remove RNA secondary structure by changing the enzyme (60).

The issue of missing the last few bases of the read, which represents the 5’-end of the transcript, is a more significant issue for those looking for single base pair resolution of transcript ends. Ligating an adaptor to the read prior to sequencing shows promise in addressing that issue (50, 61). However, these are only likely to improve recovery of the 5’-ends of transcripts, but we saw a significant amount of fragmentation at the 3’-ends that may be either incomplete transcription, 3′-degradation of transcripts, random breakage, or sequencing biases that need to be better understood.

Incomplete transcription is intriguing and may reflect the fundamental biology since (a) bacterial transcription and translation are coupled and (b) bacterial transcripts are short-lived and frequently in the process of being synthesized, since bacterial mRNAs are made at a rate of 40-80 nt/sec (62) while the average mRNA half-life is only 2-10 minutes (63). In contrast, eukaryotic RNAs have to be spliced to create mature mRNA before being exported from the nucleus and have increased stability and a longer half-life.

Some have noted the inaccuracy of ONT sequencing. There is the base inaccuracy, which should not solely be considered inaccuracy. RNA is modified with over >160 different modifications (64), and much of the inaccuracy is actually hyperaccuracy that is detecting those modifications but still trying to make assignment in four base space. It is actually the notion that RNA sequence is in four base space that is issue. That will improve as people develop base callers that expand beyond just four base space. We avoid base “inaccuracy” by using only the read mapping coordinates.

When discussing taxonomy, Stephen J. Gould emphasized that “classifications both reflect and direct our thinking” (65). Going on to say that “the way we order represents the way we think” (65). Annotation has many similarities to taxonomy, and similarly genome annotation both reflects and directs our thinking. For bacteria, annotation is protein-centric, influencing our results and ways of thinking. Historically, this is likely due to the connection between the definition of a gene and protein, but practically it also relates to the ease with which we can computationally predict proteins. However, with new experimental methods and abilities, it is time for a sea change in bacterial genome annotation. The experimental and computational methods here are inexpensive, easy, and quick, and thus they should be implemented widely. Additionally, there is a need for associated new ontology standards for describing transcripts and operons in annotation files that will better describe these features, similar to changes made in eukaryotic annotation files to accommodate alternative splicing and alternative transcripts (66). A harmonization of the two would be ideal, such that there is a standard that spans the incredible biological diversity and commonalities across the domains of life.

## Conclusions

Here we use bacterial long read transcriptome data and a new algorithm we developed to predict transcripts from this data for two strains of three diverse bacterial species including both Gram-negative and Gram-positive bacteria. Our analysis reveals a tremendous amount of transcript structural variation, transcription of RNA on both strands in some regions, overlapping transcripts, and a diversity of non-coding RNAs, which we provide as new annotation for these genomes. Bacterial transcriptional structural variation has a richness that rivals or surpasses what is seen in eukaryotes and provides a rich new set of therapeutic and diagnostic targets.

## Methods

### Bacterial cultures

Cryogenically preserved *E. coli* K12 MG1655 or E2348/69 were streaked onto an LB agar plate and placed in an incubator overnight at 37 °C. A single colony was selected to inoculate LB broth for an overnight culture. The overnight culture was diluted 1:100 in LB broth and harvested at the optical density specified in **Table 1A**. For DMEM, overnight cultures were grown in LB broth and diluted 1:100 in DMEM.

### RNA Isolation

To isolate RNA, the Qiagen RNeasy Mini Kit was used according to Qiagen RNA Protect Reagent Handbook Protocols 4 and 7 with Appendix B on-column DNase digestion (Qiagen, Hilden, Germany). The RNA was assessed with UV-Vis spectrophotometry (Denovix DS-11, Wilmington, DE), Qubit RNA HS Assay Kit (Fisher Scientific, Waltham, MA), and TapeStation RNA Screentape (Agilent, Santa Clara, CA). RNA preparations were stored at-80 °C until ready for polyadenylation and sequencing, except for the *E. coli* K12 MG1655 harvested at an optical density OD600 of 0.2. The RNA isolated from this one culture was treated four different ways. For SRR27982843, 4 μg of the freshly isolated RNA was immediately polyadenylated and then taken into library preparation and sequenced, as detailed below. The leftover polyadenyalated RNA was stored at-80 °C alongside the original RNA isolation which had been frozen without polyadenylation. Two months later, the original, unpolyadenylated RNA was thawed and polyadenylated just before library preparation and sequencing (SRR27982841). On that same day, the RNA that had been polyadenylated before being frozen was thawed and taken directly into library preparation and sequencing (SRR27982841). Four months after the original RNA isolation, the RNA that had been polyadenylated before storing at-80 °C was thawed again and polyadenylated again before library preparation and sequencing (SRR27982840).

### Oxford Nanopore Sequencing

RNA was polyadenylated with *E. coli* poly(A) polymerase (M0276S, New England Biosciences, Ipswich, Massachusetts) at 37 °C for 90 s – 30 min (Table S1) according to the manufacturer’s protocol and sequenced with the Direct RNA Sequencing kit (SQK-RNA002, Oxford Nanopore Sequencing, Oxford, UK) according to protocol version DRS_9080_v2_revR_14Aug2019. The prepared RNA library was loaded onto R9.4.1 flow cells (FLO-MIN106D) in a MinION device Mk1B (MIN-101B). Sequencing runs were terminated at 24 h. Fast5 files were basecalled using Guppy version 6.4.2 generating FASTQ files with the high accuracy model using the rna_r9.4.1_70bps_hac config file on a GPU cluster.

### Read Mapping, Transcript Prediction, and Analysis

FASTQ files were mapped to the reference genome (**Table A2**) using minimap2 (v2.24-r1122) (67)(options:-ax map-ont-t 2). Alignments were sorted and filtered with samtools (v1.11) (68) using view (option:-F 2308) and generating bam files that were merged and indexed. BED files were generated with bamToBed (v2.27.1) (69)(options:-s-c 6,4-o distinct,count) and filtered with awk to remove regions with fewer than 20 reads. The tp.py script was run in python (v.3.11.4). Statistics on regions, predicted transcripts, and other features were calculated with perl (v5.30.2). Perl (v5.30.2) was also used to merge the transcript and reference gff files and identify mRNAs, ncRNAs, and UTRs. ONT sequencing, transcript predictions, and reference CDS predictions were visualized in R (v3.6.3). E2348/69 reads from the SRA for PRJEB36845/E-MTAB-88804 and counted against the E2348/69 with the transcript predictions presented here using Salmon (v. 1.10.2) (31). Before differential expression was assessed, genes not meeting the required CPM cutoff of 5 in at least 3 samples were removed. The samples were grouped based on the treatment status, and differentially expressed genes were identified with EdgeR v3.30.3 using the quasi-likelihood negative binomial generalized log-linear model. Statistical significance was set at an FDR cutoff < 0.05 after correction with the Benjamini Hochberg method. A heatmap was drawn in R v4.2.1 using heatmap.3 of the z-score transformed log2(TPM) values for differentially expressed genes with the columns ordered based on a dendrogram generated using pvclust v2.2-0.

The full set of commands are described at: https://github.com/jdhotopp/tp.py-Direct-RNA-Sequencing-Manuscript-/tree/main (a DOI will be acquired after commands are finalized following review of the manuscript).

## Additional Files

**Table A1 – Sequencing statistics for ONT direct RNA sequencing runs with *E. coli* RNA**

**Table A2 – Data Used in the Analysis and Prediction of Transcripts**

**Figure A1 – recBCD/ptrA Transcript Predictions**

Minus-strand ONT direct RNA sequencing reads (shown as lines) are mapped from ∼2.950-2.965 Mbp in the *E. coli* K12 genome (NC_000913.3), which corresponds to a region encoding RecBCD and PtrA. Reads are sorted by their transcription stop site for *E. coli* K12 grown in rich LB media (left sorted, **A**; right sorted, **C**) and DMEM media (left sorted, **B**; right sorted, **D**). Our algorithm predicts 3 transcripts (**E**), and 7 CDSs in the reference NC_000913.3 GFF file are illustrated (**F**). While there are no ONT reads that span the entire *recBCD/ptrA* region, there is sufficient evidence to call this transcript. This is because, after removing reads wholly contained within a predicted *recBD* transcript and a *ptrA/recBD* transcript there were sufficient reads remaining to predict a transcript that spans *recBCD/ptrA*.

**Figure A2--Pore-mediated and ligase-mediated chimeras**

Chimeric sequencing reads observed in ONT direct RNA sequencing data can theoretically be generated by (A) two reads entering the pore in tandem with an open pore state that is not detected, (B) two RNA molecules being fused through ligation in vivo, in vitro during library construction, or a mapping artifact when there is a rearrangement in the genome reference, and (C) two fragments fused by ligation following adapter ligation during library construction. (D) A chimeric read from a sequencing run that shows that an open pore state (black) was missed between the first read (blue) and the second read (red). For both reads, the characteristic DNA adaptor with a lower current is observed followed by a higher plateau that is the polyA tail being sequenced. The open pore state is a spike from increased current when the pore is open between RNA molecules sequenced.

**Figure A3--Results from Stringtie, Tama, and Cupcake**

Transcript predictions resulting from Stringtie, Tama, and Cupcake are shown for *E. coli* K12 for the same region as presented in Figure 3. Plots are labeled on the right side according to the notation in the github page that fully describes how they were run with those ending in LB resulting from only the K12 LB data and those ending in DMEM resulting from only the K12 DMEM data. Stringtie is splicing focused, and since bacteria do not have splicing it is unsurprising that it could not predict transcript structures, largely yielding transcript predictions corresponding to zero-depth regions across the genome. Whether default (Tama 1) or user-defined parameters were used (Tama 2 and 3), Tama frequently over-called transcripts, particularly in regions with higher sequencing depth. With default parameters (Cupcake 1), Cupcake tends to under-call transcripts because ONT reads get filtered out of analysis due to higher degree of mismatches, and in this region no results were reported. When the parameters were adjusted to better fit ONT reads (Cupcake 2), Cupcake produced results similar to TAMA.

## Acknowledgements

This project was funded by federal funds from the National Science Foundation grant number EF 2025384 and the National Institute of Allergy and Infectious Diseases, National Institutes of Health, Department of Health and Human Services under grant numbers U19AI110820 and T32AI162579. The funding body had no role in the design of the study and collection, analysis, and interpretation of data and in writing the manuscript.

## Data availability

The ONT FASTQ file accessions for the data generated in this proposal are SRR18061005, SRR18061002, SRR27982845, SRR18061004, SRR18061003, SRR23886068, SRR27982844, SRR27982843, SRR27982842, SRR27982841, and SRR27982840.

