## Supplementary figures and images for "Deciphering Bacterial and Archaeal Transcriptional Dark Matter and Its Architectural Complexity"

### Supplementary Figures

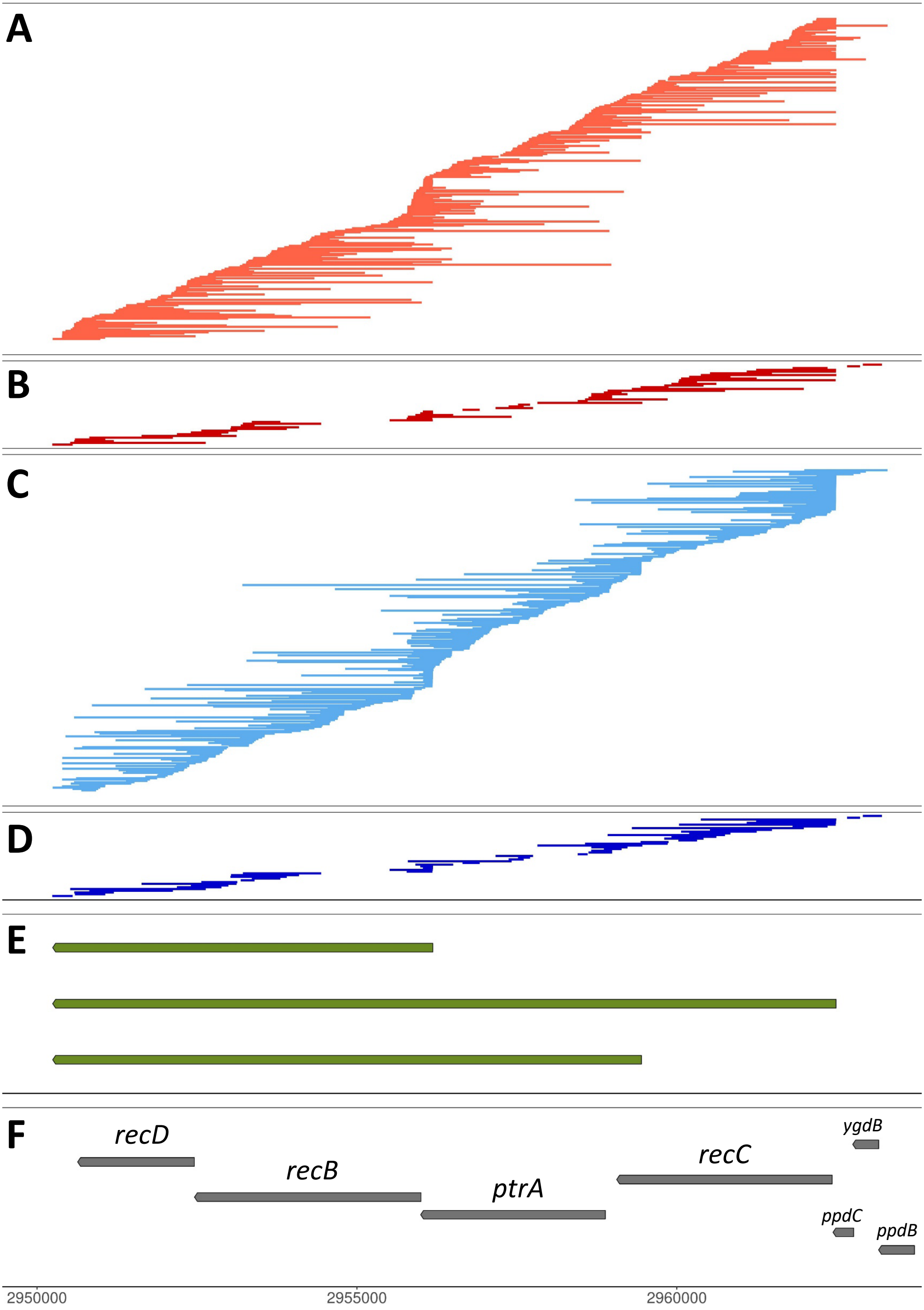

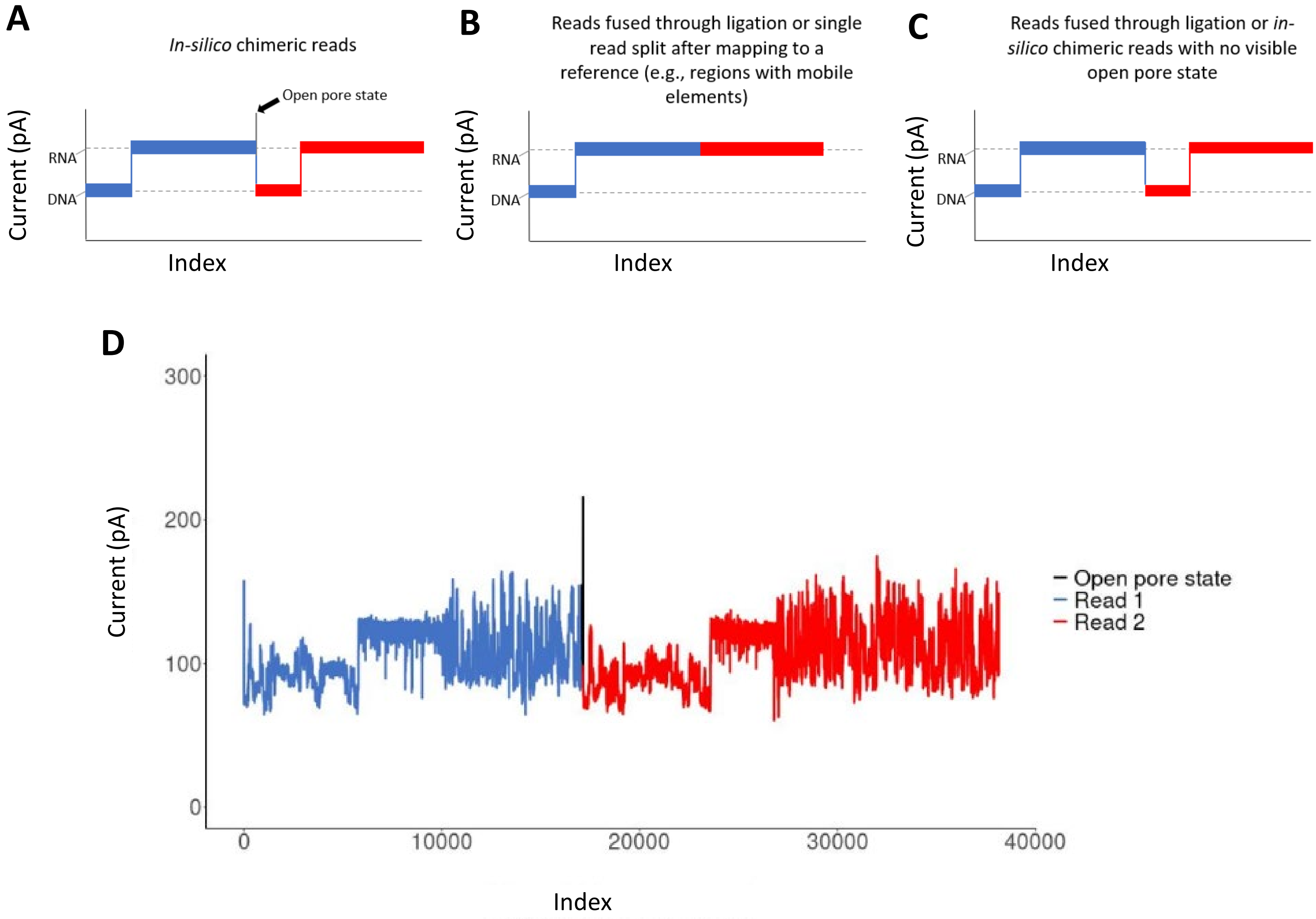

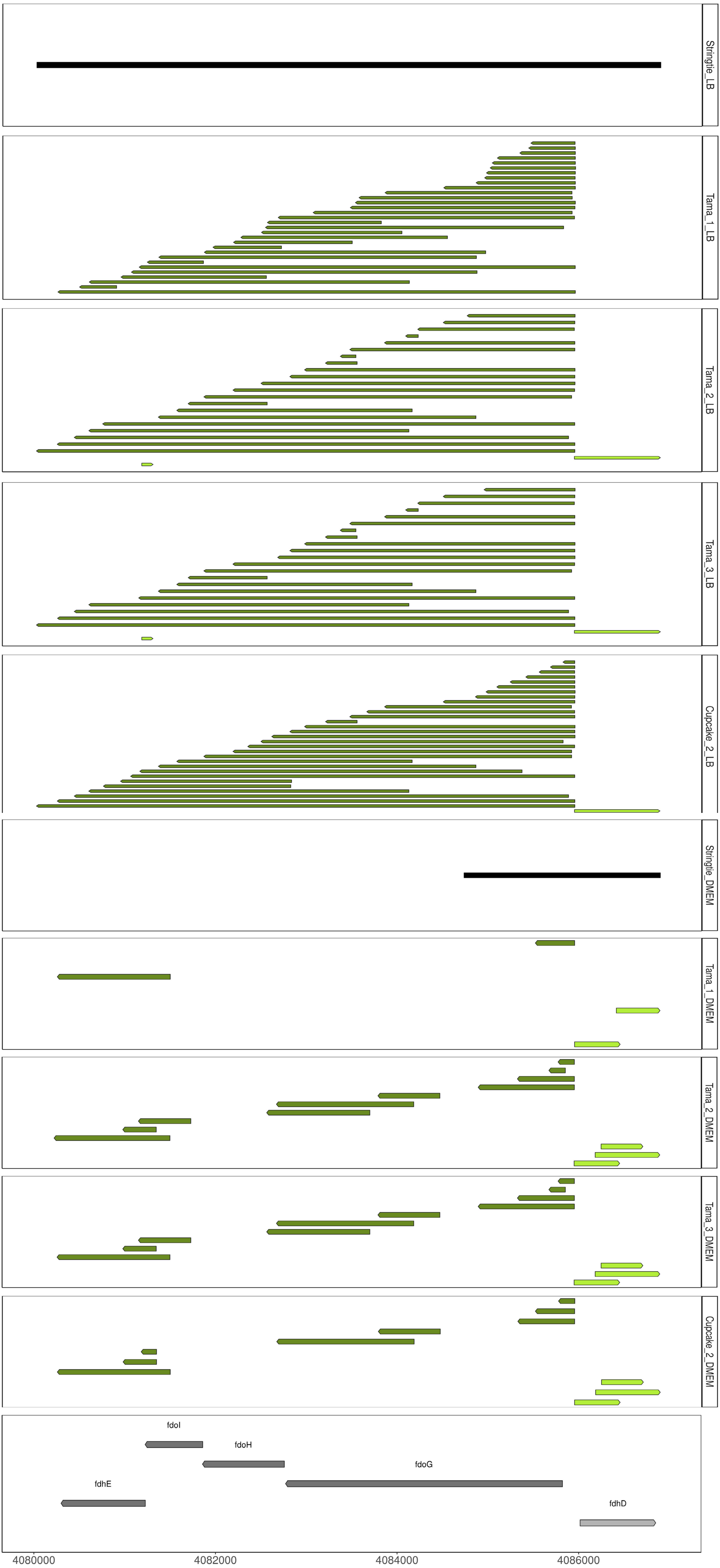
